# Explainable AI Based Coronary Heart Disease Prediction: Enhancing Model Transparency in Clinical Decision Making

**DOI:** 10.1101/2025.03.20.644456

**Authors:** Avichandra Singh Ningthoujam, Shilpa Sharma, Avishek Nandi

## Abstract

Introduction: Coronary heart disease (CHD) is still a major cause of death globally, and hence early detection and risk stratification are necessary to avoid major cardiovascular events. The present study uses clinical and demographic characteristics to compare the predictive accuracy of eight machine learning models for CHD diagnosis. It also investigates the contribution and direction of influence of the most important features in the models to improve interpretability.

**Methods:** We contrasted the predictive accuracy of eight different machine learning models for CHD classification. The work identifies the most important features from the top-performing models and applies SHapley Additive exPlanations (SHAP) and Local Interpretable Model-Agnostic Explanations (LIME) to gain insight into how every feature affects the model’s prediction. These interpretive methods assist in displaying the direction and amount of feature contributions to allow transparency in AI-based CHD risk prediction.

**Results:** XGboost and Random Forest achieved the highest testing accuracies 0.839 and 0.805, with training accuracies of 0.901 and 0.957 respectively showing ideal model of XGboost and significant overfitting for random forest model. ECG-associated features, such as resting ECG and old peak (ST depression through workout), also place favorably, supporting the significance of cardiac electrical activity in diagnosis. ST slope has the highest impact, followed by Chest pain type and old peak, which increase the likelihood of heart disease with their high values contributing positively to the prediction. Resting bps, sex, and fasting blood sugar have lower impacts on the model’s predictions.

**Conclusions:** In conclusion, machine learning models, particularly XGboost and random forest, show substantial predictive accuracy for coronary heart disease, with testing AUROCs of 0.885. Feature importance, SHAP and LIME analysis highlight the critical role of ECG-derived metrics like ST slope, chest pain type, and resting ecp while traditional risk factors such as cholesterol, resting bps, and fasting blood sugar have less influence.

## Introduction

Coronary Heart Disease (CHD) remains one of the foremost origins of sickness and mortality globally, placing an enormous strain on public health systems [1]. Early diagnosis and accurate risk stratification are crucial to managing CHD, enabling timely interventions and more effective treatments [2]. By tradition, clinical measurements and demographic records have been utilized to classify at-risk people. Conversely, with the introduction of machine learning, there is currently a break to enhance analytical precision by adding numerous data points into full demonstrates [3] [4]. Recent research has shown the potential of various machine learning (ML) algorithms in forecasting CHD using clinical and demographic features [5], [6]. In this study, we focus on a set of attributes that are routinely collected in clinical settings: age, sex, chest pain type, resting blood pressure (bps), cholesterol, fasting blood sugar, resting electrocardiographic (ECG) results, maximum heart rate achieved, exercise-induced angina, old peak (an indicator of ST depression), and ST slope [7], [8].

These features specify a complete picture of a patient’s cardiovascular condition and suggest diagnostic perceptions of the primary pathophysiology of CHD [9]. For example, demographic components such as age and sex are familiar judges of CHD, with aged persons and males in general demonstrating a greater probability [10]. Similarly, a patient’s chest pain can show the importance of cardiac issues [11]. The typical concern specifies clinical measures such as resting blood pressure, cholesterol levels, and fasting blood sugar, demonstrating the patient’s cardiac and metabolic situation [12]. Moreover, diagnostic tests, including resting ECG and exercise metrics like max heart rate and exercise-induced angina, help assess cardiac function and stress response [13], [14]. Additionally, parameters like old peak and ST slope provide quantifiable measures of ST-segment changes during exercise, which are critical for identifying myocardial ischemia [15], [16]. Despite the established significance of these features, there remains considerable variability in how well current models predict CHD. Recent advances in ML suggest the ability to address variability by effectively integrating and analyzing multifaceted data to yield more precise predictions [17]. Our work compares different ML procedures to develop and validate projecting models that harness these clinical and demographic features [18]. We try to find the highest effective algorithmic approach for early CHD prediction, facilitating improved clinical decision-making and personalized patient maintenance. CHD remains a leading cause of mortality worldwide, prompting extensive research into its prediction, diagnosis, and management using computational and clinical approaches. Machine learning (ML) and deep learning (DL) approaches have improved early detection and prognosis accuracy [19]. Several studies have leveraged traditional machine learning algorithms, such as logistic regression (LR) and support vector machines (SVM) for CHD prediction. Li et al. [20] developed an ML-based predictive model that integrated clinical risk factors and electrocardiogram (ECG) features, achieving a sensitivity of over 85%. Similarly, Zhang et al. [21] proposed an ensemble learning approach combining multiple classifiers to enhance prediction performance. Deep learning techniques, particularly convolutional neural networks (CNNs) and recurrent neural networks (RNNs) have presented favorable outcomes in CHD detection using medical imaging and ECG signals. Wang et al. [22] implemented a CNN-based model for automated CHD classification from echocardiographic images, achieving state-of-the-art accuracy. Additionally, hybrid models integrating CNNs with long short-term memory (LSTM) networks have revealed exceptional execution in time-series data analysis [23]. Recent studies have also discovered the role of explainable AI (XAI) in CHD diagnosis to improve transparency and interpretability. A study by Kumar et al. [24] incorporated SHAP (Shapley Additive Explanations) values to detect the highly effective risk factors promoting to CHD predictions, aiding clinicians in decision-making. Despite these developments, challenges such as imbalanced datasets, the need for normalized symptomatic measures, and real-time application restraints keep at [25]. The potential studies should expand the model overview, incorporate multi-modal data resources, and develop more explainable models to improve clinical adoption. Rationale and knowledge gap: Despite advances in CHD treatment, early detection remains a challenge. Machine learning can potentially improve prediction accuracy, but its comparative effectiveness across different models is not well-studied [26]. Most CHD prediction studies focus on a single machine-learning model. However, limited research compares multiple ML algorithms using diverse clinical and demographic features to determine the best-performing model [27]. Objective: This research aims to develop and evaluate ML versions for predicting coronary heart disease (CHD) using clinical and demographic attributes. Additionally, it analyzes and visualizes the impact of different features on the predictive models, highlighting their significance and directional influence. [28]. Specifically, this study judges the predictive performance of eight machine learning algorithms for CHD classification. Recognize the most precise features and consistent model for early CHD risk evaluation [29]. Analyze the impact of key clinical and demographic features on CHD prediction. Specify perceptions into the possible clinical application of ML models for early intervention and improved patient outcomes.

## Materials and methods

### 0.1 Study design

This study follows a retrospective observational design, utilizing a publicly available dataset to develop and evaluate ML models for coronary heart disease (CHD) prediction [30]. The research focuses on applying supervised learning techniques to classify individuals based on their CHD risk using clinical and demographic attributes [31] and visualizes the impact of different features on the predictive models, highlighting their significance and directional influence.

### 0.2 Data Collection

The Cleveland Clinic Heart Disease Dataset is obtained from Kaggle with 1048 datasets, 520 heart diseases, and 528 no heart diseases [32], [33]. The dataset has 11 features and is as follows: age: age in years, sex: sex (1 = male; 0 = female), cp: chest pain type: Value 1: typical angina, Value 2: atypical angina, Value 3: non-anginal pain and Value 4: asymptomatic, fbs: fasting blood sugar *>* 120 mg/dl (1 = true; 0 = false), Cholesterol level, fasting blood sugar: value 1 for true and 0 for false, restecg: resting electrocardiographic results: Value 0: normal, Value 1: having ST-T wave abnormality (T wave inversions and/or ST elevation or depression of *>* 0.05 mV), Value 2: showing probable or definite left ventricular hypertrophy by Estes’ criteria, thalach: maximum heart rate achieved, exang: exercise induced angina (1 = yes; 0 = no), oldpeak: ST depression induced by exercise relative to rest, slope: the slope of the ST peak exercise segment: Value 0: horizontal, upsloping, Value 1: upsloping, Value 2: flat, Value 3: downsloping and finally Target 1 for heart disease and 0 for no heart disease [34], [35]. We apply various data pre-processing steps, including the removal of highly correlated attributes. The data set is then split into training and testing sets in an 80:20 ratio to train the models.

### 0.3 Statistical analysis

For statistical analysis, present count, mean, and standard deviation for continuous features such as age, cholesterol, and blood pressure and Frequency distribution for categorical variables (e.g., sex, chest pain type) [36] [37]. Statistical analysis and machine learning pipelines should leverage key Python libraries such as tableone, Pandas and NumPy, Matplotlib and Seaborn, Scikit-learn, and SciPy and Statsmodels using Python version 3.12.3 [38].

### 0.4 Machine Learning Based Algorithms

For model evaluation, we have conducted eight ML algorithms, including tree-and ensemble-based models: eXtreme Gradient Boosting (XGBoost), random forest (RF), and Decision tree; a distance-based model: support vector machine (SVM); Logistic Regression, K-NN, Naive Bayes as well as Deep learning model [39], [40], [41].

### 0.5 Model Performance Evaluation and interpretation

We present accuracy and Area Under the Receiver Operating Characteristic (AUROC) to assess model performance and measure the model’s capability to differentiate between CHD vs non-CHD classes; finally, 95% Confidence Interval (CI) for the AUROC Bootstrapping method to compute the confidence interval for AUROC and assess model robustness [42], [43]]. To understand the ML models, we applied the Shapley additive extension (SHAP) and Local Interpretable Model-Agnostic Explanations (LIME) values to clarify the way and intensity of the picked attributes in the ultimate model [44].

## Results

In Table 1 it provides a detailed comparison of patient characteristics between those with and without heart disease, along with statistical measures (SMD and P-values) to assess differences. The analysis of patient characteristics highlights several key factors associated with heart disease. Patients with heart disease are inclined to be older (54.8 vs. 51.9 years, P*<*0.001) and have a higher proportion of males (78.5% vs. 68.6%, P*<*0.001). Chest pain type is a strong differentiator, with asymptomatic pain being more prevalent in heart disease cases (55.8% vs. 19.5%, P*<*0.001). Clinical markers such as resting ECG abnormalities, lower maximum heart rate, and higher resting blood pressure also show statistically significant differences. Exercise-induced angina (50% vs. 23.9%, P*<*0.001) and ST segment slope abnormalities (SMD=1.41, P*<*0.001) are among the strongest predictors. At a similar interval, cholesterol amounts do not demonstrate a statistically significant difference (P=0.069), indicating restricted analytical meaning in this dataset.

**Table 1.** Baseline Characteristics of Study Participants.

| Variable | Categories | Overall (n=1048) | No HD (n=528) | HD (n=520) | P-Value |
| --- | --- | --- | --- | --- | --- |
| Age, mean (SD) |  | 53.3 (9.4) | 51.9 (9.4) | 54.8 (9.2) | <0.001 |
| Sex, n (%) | Female | 278 (26.5) | 166 (31.4) | 112 (21.5) | <0.001 |
|  | Male | 770 (73.5) | 362 (68.6) | 408 (78.5) |  |
| Chest Pain Type, n (%) | Typical Angina | 184 (17.6) | 130 (24.6) | 54 (10.4) | <0.001 |
|  | Atypical Angina | 216 (20.6) | 154 (29.2) | 62 (11.9) |  |
|  | Non-anginal Pain | 255 (24.3) | 141 (26.7) | 114 (21.9) |  |
|  | Asymptomatic | 393 (37.5) | 103 (19.5) | 290 (55.8) |  |
| Resting BPS, mean (SD) |  | 132.6 (17.4) | 131.3 (16.9) | 134.0 (17.7) | 0.012 |
| Cholesterol, mean (SD) |  | 245.2 (57.1) | 242.0 (54.1) | 248.4 (59.8) | 0.069 |
| Fasting Blood Sugar (>120 mg/dl) n (%) | FALSE | 878 (83.8) | 463 (87.7) | 415 (79.8) | 0.001 |
|  | TRUE | 170 (16.2) | 65 (12.3) | 105 (20.2) |  |
| Resting ECG, n (%) | Normal | 592 (56.5) | 336 (63.6) | 256 (49.2) | <0.001 |
|  | Abnormality | 276 (26.3) | 108 (20.5) | 168 (32.3) |  |
|  | Hypertrophy | 180 (17.2) | 84 (15.9) | 96 (18.5) |  |
| Max Heart Rate, mean (SD) |  | 142.9 (24.4) | 146.5 (23.4) | 139.3 (25.0) | <0.001 |
| Exercise Induced Angina n (%) | No | 662 (63.2) | 402 (76.1) | 260 (50.0) | <0.001 |
|  | Yes | 386 (36.8) | 126 (23.9) | 260 (50.0) |  |
| Oldpeak, mean (SD) |  | 0.9 (1.1) | 0.7 (1.0) | 1.2 (1.1) | <0.001 |
| ST Slope, n (%) | Horizontal | 22 (2.1) | 12 (2.3) | 10 (1.9) | <0.001 |
|  | Upsloping | 489 (46.7) | 395 (74.8) | 94 (18.1) |  |
|  | Flat | 494 (47.1) | 110 (20.8) | 384 (73.8) |  |
|  | Downsloping | 43 (4.1) | 11 (2.1) | 32 (6.2) |  |
| Target, n (%) | No Heart Disease | 528 (50.4) | 528 (100.0) | — | <0.001 |
|  | Heart Disease | 520 (49.6) | — | 520 (100.0) |  |

Data are presented as number (percentage) for categorical variables and mean (SD standard deviation) for continuous variables. HD=Heart Disease

Combining Standardized Mean Difference (SMD) and P-values provides a comprehensive understanding of the data. Large SMD values indicate meaningful differences in features such as chest pain type and ST slope, reinforcing their importance in predictive modelling [45]. Even when the SMD is small, a significant P-value confirms a variable’s relevance, as seen with age and fasting blood sugar. The dataset is nearly balanced (520 heart disease vs. 528 no heart disease cases), reducing model bias.

Table 2 compares various machine learning models’ training and testing accuracies, highlighting their generalization capabilities and potential overfitting. Models like Random Forest (95.7% training vs. 80.5% testing) and k-NN (85.3% training vs. 75.2% testing) show significant gaps between training and testing performance, suggesting overfitting due to high model complexity. In contrast, XGBoost achieves the best testing accuracy (83.9%) with a smaller gap (90.1% training), indicating strong generalization through built-in regularization. The Deep Learning Model also balances performance (80.0% training vs. 80.0% testing), likely due to effective techniques like dropout or early stopping. Logistic Regression somewhat breaks on testing (79.5%) evaluated to training (77.8%), maybe as its modesty avoids overfitting. At the same time, Naive Bayes fails (71.9% testing) due to its feature-independent status belief, which may not support the dataset’s arrangement.

**Table 2.** Different Machine Learning Models Performance Evaluation.

| Models | Training Accuracy | Testing Accuracy |
| --- | --- | --- |
| Decision Tree | 0.785 | 0.786 |
| Logistic Regression | 0.778 | 0.795 |
| Naive Bayes Model | 0.728 | 0.719 |
| k-NN Model | 0.853 | 0.752 |
| SVM Model | 0.833 | 0.791 |
| Random Forest Model | 0.957 | 0.805 |
| XGBoost | 0.901 | 0.839 |
| Deep Learning Model | 0.7995 | 0.800 |

Table notes: The table displays the training and testing accuracies for various machine learning models evaluated in the study.

Table 3 compares the performance of different ML models using the AUROC for training and testing datasets, along with 95% confidence intervals (CI). The Decision Tree and Logistic Regression models show comparable testing AUROCs (0.784), but Logistic Regression has a higher training AUROC (0.835 vs. 0.785), suggesting it may generalize better despite similar testing performance. The Naive Bayes model is notable for its slight improvement in testing AUROC (0.805) compared to training (0.800), indicating robust generalization. More complex models like k-NN, SVM, Random Forest, and XGBoost exhibit high training AUROCs (0.931–0.996), but their testing scores drop significantly (0.836–0.885), highlighting potential overfitting. The Deep Learning Model balances moderate training performance (0.870) with a competitive testing AUROC (0.853), reflecting a trade-off between complexity and generalizability. The Random Forest and XGBoost models achieve the highest testing AUROCs (0.885) and 0.996 and 0.967 for training respectively, suggesting they capture complex patterns while maintaining reasonable generalization which is shown in Fig 1 for testing and Fig 2 for training datasets. The k-NN and SVM models also show strong testing performance (0.836–0.844) but with wider confidence intervals, indicating variability—logistic Regression testing AUROC (0.784) overlays by its training CI, indicating constant performance. The 95% CIs for testing AUROCs frequently overlay across models, signifying statistical doubt in place them ultimately. The optimal may differ on balancing performance, interpretability, and computational cost for real-world applications. Models like Random Forest or XGBoost might be preferred for high accuracy. At the same time, Logistic Regression or Naive Bayes could be chosen for simplicity and efficiency if slight performance trade-offs are acceptable.

**Fig 1.**
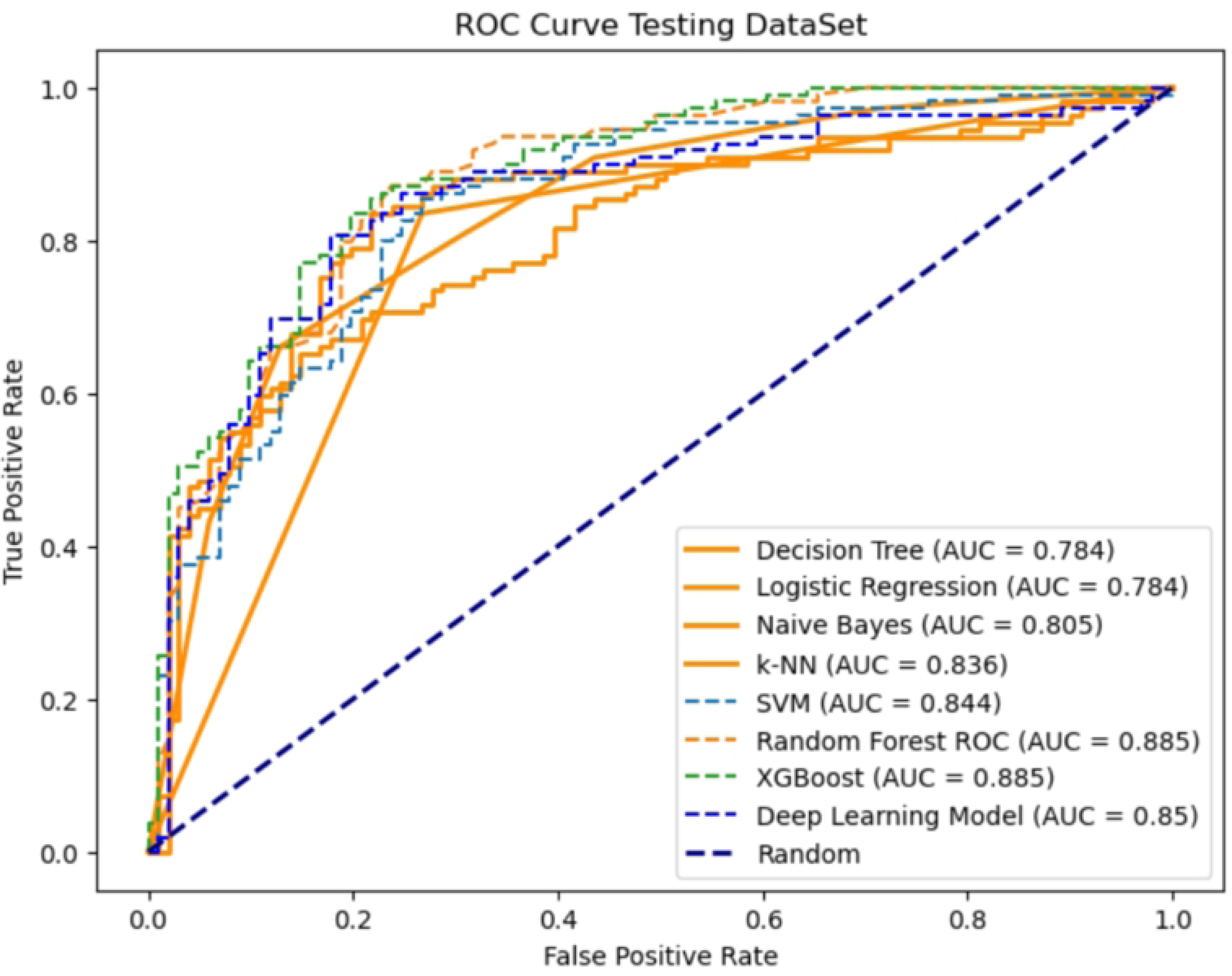
Area Under the Receiver Operating Characteristic Curve (AUROC) for Testing dataset.

**Fig 2.**
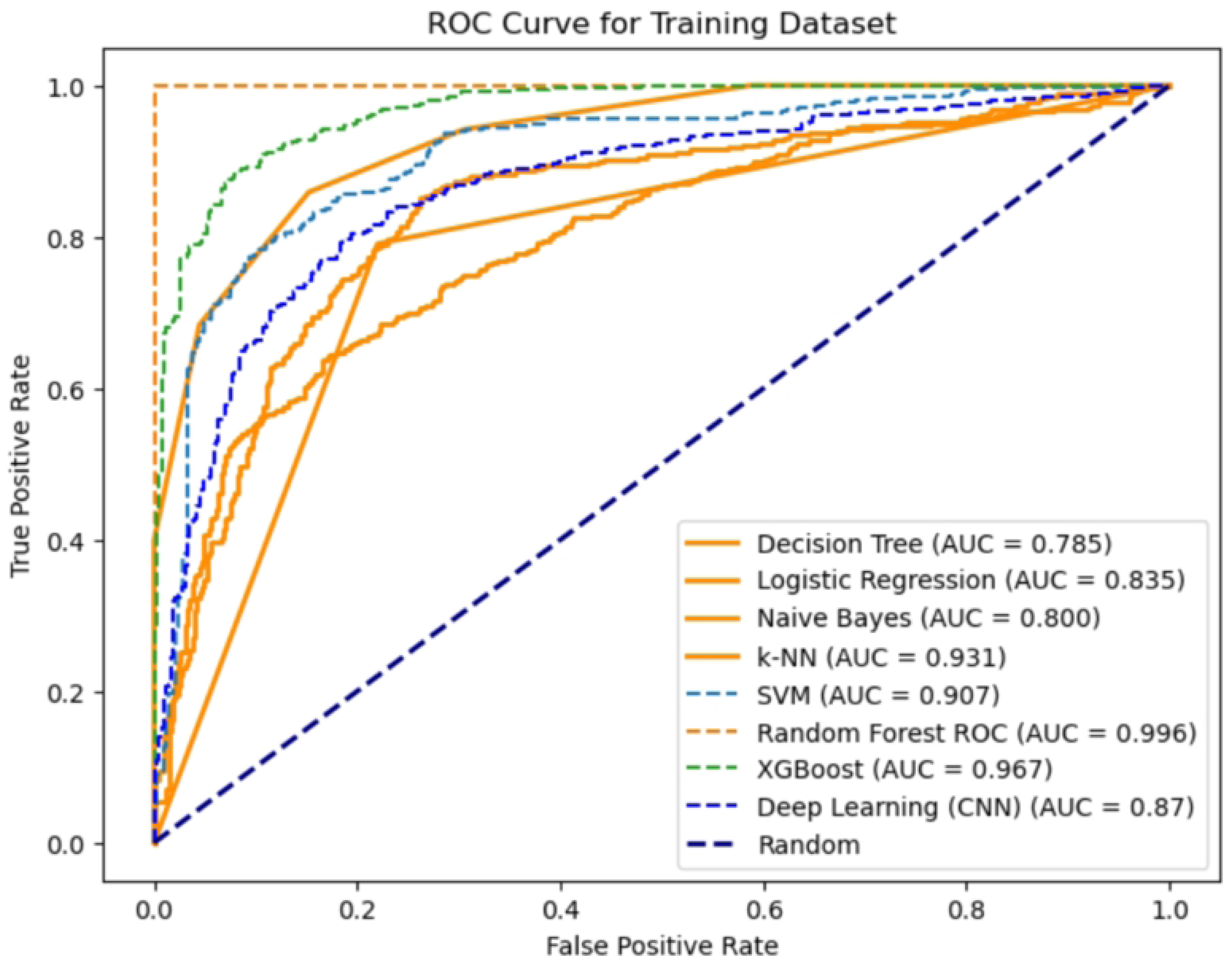
Area Under the Receiver Operating Characteristic Curve (AUROC) for Training dataset.

**Table 3.**
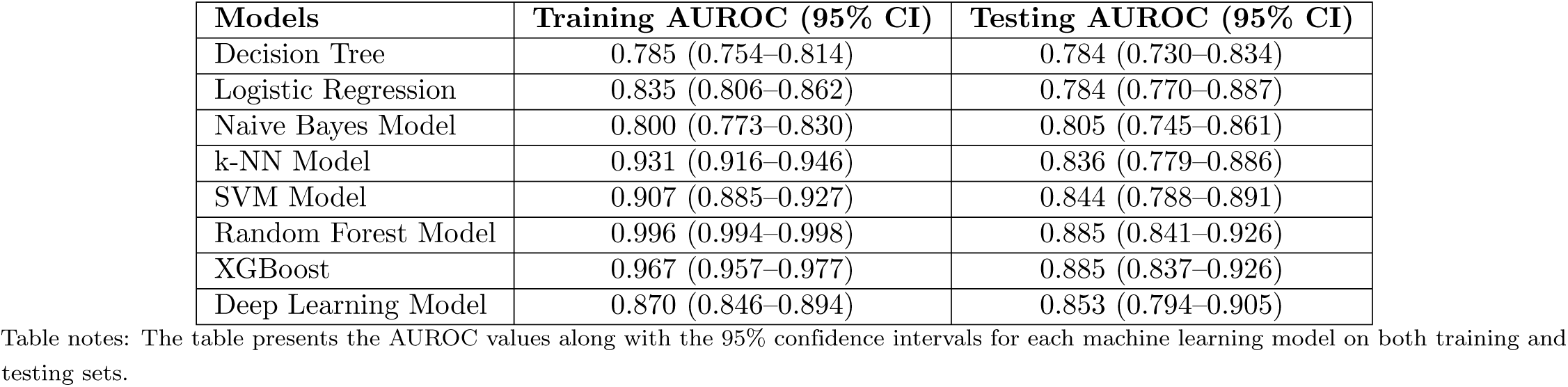
Area Under the Receiver Operating Characteristic Curve (AUROC) for training and testing.

| Models | Training AUROC (95% CI) | Testing AUROC (95% CI) |
| --- | --- | --- |
| Decision Tree | 0.785 (0.754–0.814) | 0.784 (0.730–0.834) |
| Logistic Regression | 0.835 (0.806–0.862) | 0.784 (0.770–0.887) |
| Naive Bayes Model | 0.800 (0.773–0.830) | 0.805 (0.745–0.861) |
| k-NN Model | 0.931 (0.916–0.946) | 0.836 (0.779–0.886) |
| SVM Model | 0.907 (0.885–0.927) | 0.844 (0.788–0.891) |
| Random Forest Model | 0.996 (0.994–0.998) | 0.885 (0.841–0.926) |
| XGBoost | 0.967 (0.957–0.977) | 0.885 (0.837–0.926) |
| Deep Learning Model | 0.870 (0.846–0.894) | 0.853 (0.794–0.905) |

Table notes: The table presents the AUROC values along with the 95% confidence intervals for each machine learning model on both training and testing sets.

### 0.6 Features Important

The important feature scores from the XGBoost model reveal which factors most strongly influence predictions of heart disease, offering insights into clinical relevance and model behavior. The ST slope (a measure of ECG changes during exercise) is the most critical feature (importance = 0.507), suggesting it is a key indicator of myocardial ischemia or heart stress which is shown in Fig 3. This is associated with health expertise, as unusual ST sections are frequently associated with coronary artery disease [46], [47]]. Additional ECG-associated features, such as resting ECG and old peak (ST depression through workout), also place favorably, supporting the significance of cardiac electrical activity in diagnosis. Symptoms like chest pain type and exercise angina (chest pain triggered by physical exertion) are essential, reflecting their role in clinical assessments of heart disease.

**Fig 3.**
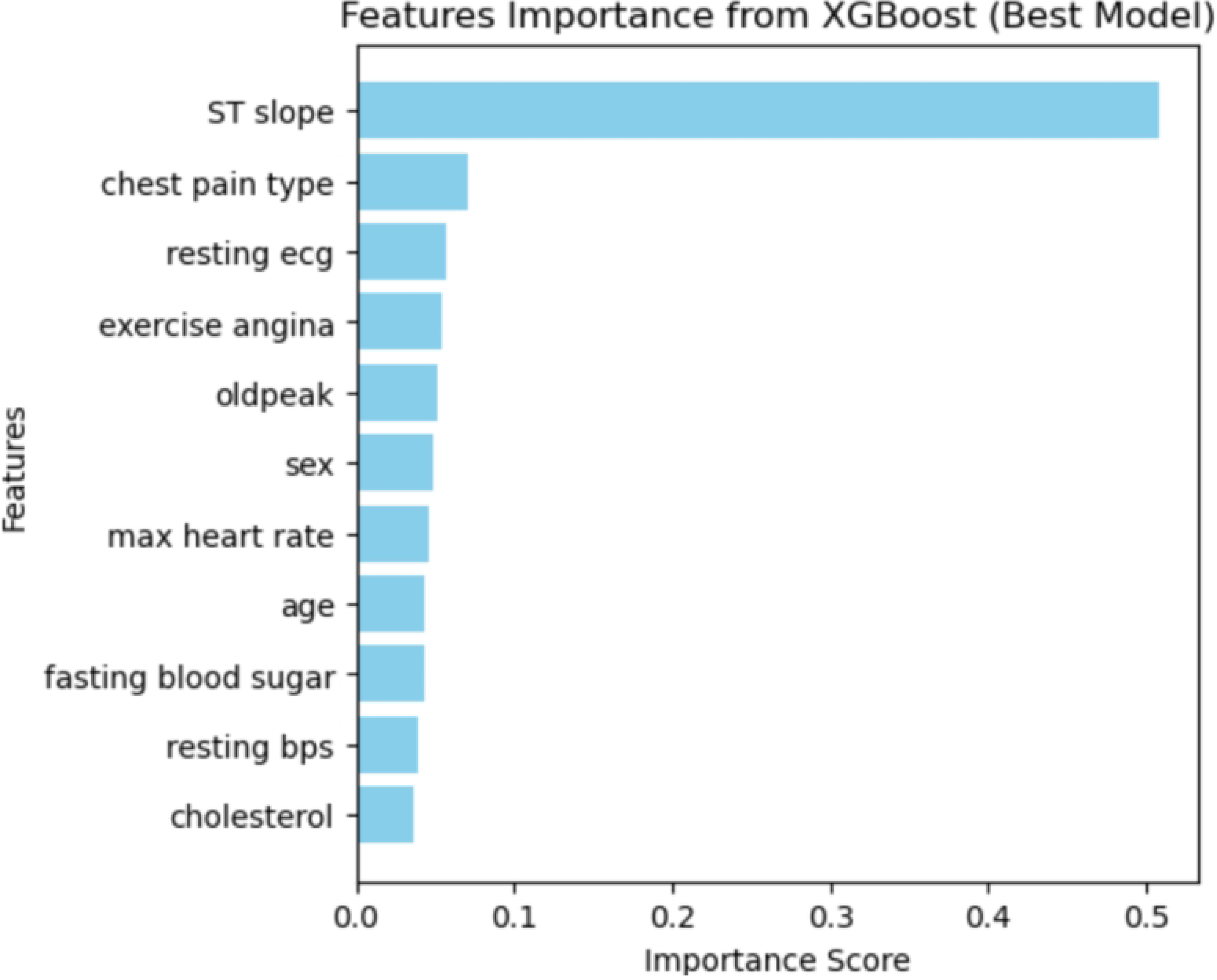
Important Features from Best Performance XGBoost Model.

**Fig 4.**
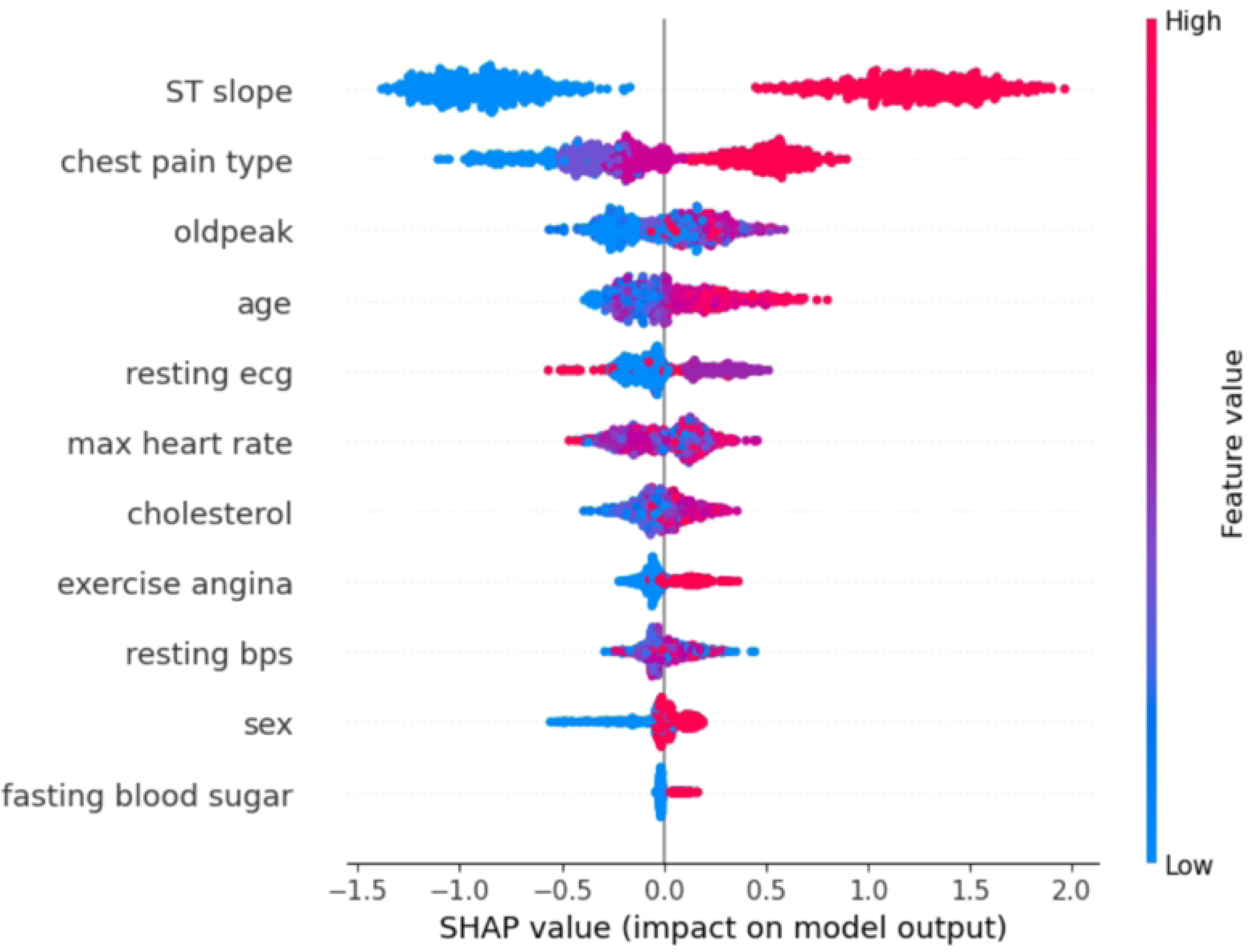
Shapley Additive exPlanations (SHAP) summary plot to explain the feature importance obtained by the XGBoost algorithm.

Demographic factors (sex, age) and traditional cardiovascular risk factors (resting blood pressure, cholesterol, fasting blood sugar) have moderate influence, highlighting that while they contribute, they are less discriminatory in this model compared to direct physiological markers.

### 0.7 SHAP evaluation

Features with higher feature value (red) and positive SHAP value (on the right side) illustrated in the figure show a positive association, while features with higher feature value (blue) and negative SHAP value (on the left side) show a negative association) [48], [49]. The supremacy of ECG-derived attributes (ST slope, old peak) suggests the algorithm focuses on objective clinical assessment of self-reported symptoms or baseline demographics. This advises that involvements or datasets missing specified cardiac testing (e.g., resting or exercise ECG) could reduce the algorithm’s performance. The relatively slighter concern of cholesterol and blood pressure opposes their value in traditional probability evaluations, feasibly because these features are more common in the general population or do not concern ECG reports.

This points out the necessity for high-quality ECG reports and rational assessment for reasonable application which is shown in Fig 5. Though the algorithms influence clinical signs effectively, they can misjudge lifestyle-associated threat features that are actionable for stopping. Estimating these readings with clinical shows could enhance algorithms’ working and real-world applicability.

**Fig 5.**
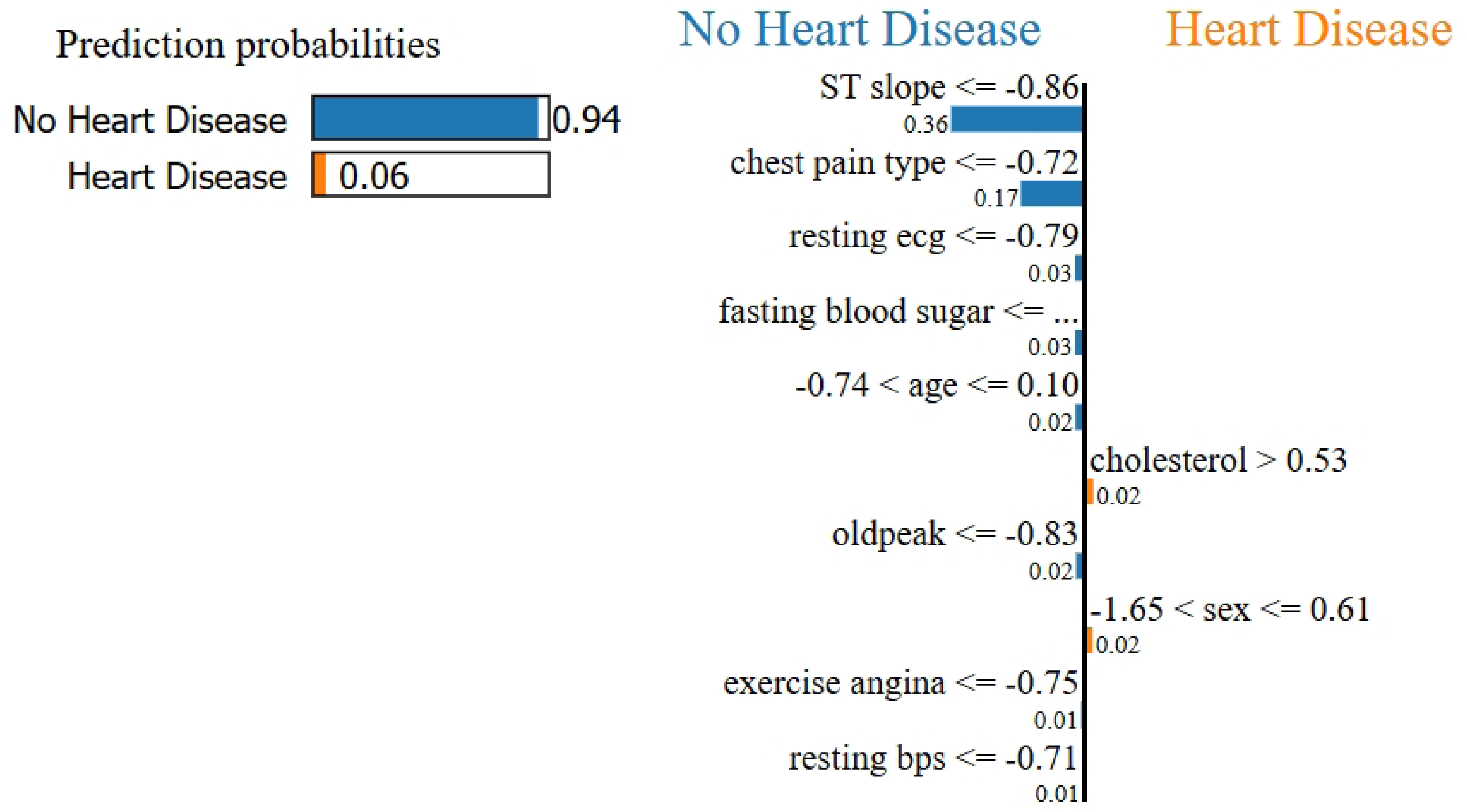
Local Interpretable Model-Agnostic Explanations (LIME) Explaination for XGBoost.

### 0.8 Local Interpretable Model-Agnostic Explanations (LIME)

LIME is a technique designed to explain individual estimates of any ML model by estimating it closely with an understandable model. By agitating the input data and perceiving the matching variations in the output, LIME figures a more straightforward, explainable model that imitators the presentation of the authoritarian model in the area of the clear presumption, thus improving the transparency and reliability of ML applications. In Fig 5 the blue color indicates the features favoring no heart disease, and the orange color favors heart disease. ST slope *≤ −*0.86 is the strongest factor to recommend the No Heart Disease result and has the strongest contribution with a value of 0.36. Other important factors are chest pain type *≤ −*0.72 (0.17), oldpeak *≤ −*0.83 (0.03), age between *−*0.74 and *−*0.10 (0.03), and resting ECG *≤ −*0.79 (0.03). Other factors like exercise angina, fasting blood sugar, and maximum heart rate also contribute to this prediction to a lesser extent.

In contrast, the characteristics driving the model towards Heart Disease are weaker in their effects. The most prominent is sex with a score of between -1.65 and 0.61 contributing 0.04 to the heart disease likelihood. Cholesterol ¿ 0.53 adds another 0.02. As the impact of these risk factors is modest in comparison with the stronger drivers for No Heart Disease, the model ends up giving a weak (5%) prediction of heart disease. The model’s prediction of ’No Heart Disease’ is primarily influenced by the ’ST Slope’ and ’Chest Pain Type’ features, as they have the highest weights. Other features like ’Sex,’ ’Oldpeak,’ and ’Age’ also contribute to a lesser extent. In clinical terms, these features relate to the ST Slope: The peak exercise ST segment’s slope indicates heart stress levels. Chest Pain Type: The nature of chest pain experienced can indicate heart conditions. Old peak: ST depression induced by exercise relative to rest, measuring heart stress. Resting ECG: Results from the resting electrocardiogram show the heart’s electrical activity. Exercise Angina: Occurrence of angina during exercise, indicating potential heart issues. Max Heart Rate: The maximum heart rate achieved during exercise is relevant to heart health.

## Discussion

The outcome of the study discloses that machine learning models can efficiently predict coronary heart disease (CHD) using clinical and demographic attributes. Between the eight ML models, ensemble-based methods such as XGBoost and Random Forest demonstrated higher analytical performance, attaining higher AUROC results and classification accuracy than traditional models like Logistic Regression and Decision Tree. The deep learning model, despite its complexity, showed competitive performance but required more extensive tuning to optimize results. These outcomes are associated with prior results that highlight the improvements of ensemble learning in conducting non-linear associations and difficult feature relations in medical datasets. In addition, feature importance investigation highlighted key risk features such as age, cholesterol levels, and blood pressure, strengthening their clinical impact in CHD risk assessment. Key Predictors of Heart Disease: Type of chest pain (asymptomatic), ST slope abnormalities, exercise-induced angina, and old peak exhibit high SMD values and highly significant p-values, presenting them as great signs of heart disease. Features such as age, sex, fasting blood sugar, and ECG abnormalities also act as major discriminators. Cholesterol (P=0.069) does not show statistical significance, suggesting it may not be a strong standalone predictor. In machine learning models, variables with large SMDs, such as ST slope and chest pain type, are likely to be strong predictors. P-values should be used to prioritize features for clinical decision-making [50], [51].

While age has a normal SMD (0.32) with a P-value (*<*0.001), indicating its role as an ideal predictor. On the contrary, cholesterol’s absence of significance (P=0.069) suggests it requires further study. The results emphasize trade-offs between complexity and generalization. XGBoost is the best test data performer, making it perfect for functions selecting accuracy. As compared, more straightforward algorithms such as Logistic Regression or Decision Trees (78.5% testing) advance appropriate interpretability with practical performance. Overfitting in models like Random Forest or k-NN could be mitigated by tuning hyperparameters (e.g., reducing tree depth, increasing k) or collecting more data. Meanwhile, SVM (79.1% testing) shows moderate overfitting, suggesting room for optimization via kernel selection or regularization. The weak performance of Naive Bayes underlines the importance of proving algorithm rules besides data characteristics. Generally, the optimal varies on the project: focus on XGBoost or deep learning for accuracy, näıve algorithms for simplicity, and focus overfitting during parameter tuning or data augmentation. The study demonstrates that machine learning models can improve early risk assessment and stratification, aiding in proactive patient management. Machine learning models, especially sophisticated ones like deep learning and ensemble techniques, tend to be black boxes, and their decision-making is difficult to interpret. This lack of interpretability presents strong challenges, particularly in high-stakes applications such as healthcare, where knowing how a model arrives at a diagnosis is necessary for trust and verification. This LIME (Local Interpretable Model-agnostic Explanations) plot illustrates the prediction of an XGBoost machine learning model for heart disease classification. The prediction probabilities show that the model overwhelmingly predicts the No Heart Disease outcome with a 96% probability, and the probability of Heart Disease is merely 4%. This implies that, the model predicts with certainty the lack of heart disease. In this instance, most of the factors point towards the absence of heart disease, and a high-confidence prediction is made for No Heart Disease.

### 0.9 Comparison with similar research

The decisions of our study are associated with previous studies on machine learning-based CHD prediction, supporting the efficiency of ensemble methods and deep learning for cardiovascular risk judgment [52]. Earlier findings have shown that XGBoost and Random Forest constantly accomplish superior predictive accuracy compared to traditional models like Logistic Regression and Decision Tree [53], [54]. Similarly, our study confirms that these ensemble techniques outperform simpler models due to their ability to handle complex, non-linear relationships in medical data. Compared to previous studies that used smaller datasets with limited feature selection, our research comprehensively evaluates eight ML models and emphasizes the importance of diverse clinical attributes.

## Conclusion

The results of this study indicate that ensemble-based machine learning models, particularly XGBoost and Random Forest, achieved the highest predictive accuracy and AUROC scores. This superior performance can be attributed to their ability to handle non-linear relationships and interactions between multiple risk factors. These findings align with previous studies that emphasize the robustness of ensemble methods in medical data classification. The feature importance analysis in this study highlights key clinical and demographic attributes that significantly contribute to coronary heart disease (CHD) prediction. Among these, ST Slope, chest pain type, resting ECG, and exercise angina have emerged as the most influential factors, and features such as cholesterol, resting BPS, fasting blood sugar, and age are the least influential factors, which align with established cardiovascular risk assessment models. This study finds that machine learning improves CHD prediction accuracy, and AI can enhance early diagnosis and patient outcomes. Healthcare providers should integrate ML-based predictive models into routine screening for high-risk patients. This investigation reveals that machine learning algorithms can efficiently determine coronary heart disease (CHD) with clinical and demographic characteristics. Between the eight models calculated, ensemble-based approaches (XGBoost and Random Forest) outperformed traditional methods in terms of accuracy and AUROC, showing their strength in managing complicated, non-linear associations. The deep learning model also showed competitive performance, but its lack of interpretability remains challenging for clinical adoption. Feature importance analysis identified: ST slope has the highest impact, with high values significantly increasing the likelihood of heart disease. Chest pain type and old peak also show substantial influence, with their high values contributing positively to the prediction. Age and resting ecg have moderate impacts, with higher values generally increasing the risk. Features like max heart rate, cholesterol, and exercise angina show varying impacts depending on their values. Resting bps, sex, and fasting blood sugar have lower impacts on the model’s predictions. Future work should focus on expanding the dataset to more diverse populations and incorporating explainable AI techniques to improve transparency and clinical integration. Integrating machine learning into CHD prediction can ultimately enhance early diagnosis, risk stratification, and personalized treatment, improving patient outcomes.

## Supporting information

## Acknowledgments

The authors acknowledge Andras Janosi, M.D., Hungarian Institute of Cardiology. Budapest. William Steinbrunn, M.D., University Hospital, Zurich, Switzerland. Matthias Pfisterer, M.D., University Hospital, Basel, Switzerland and Robert Detrano, M.D., Ph.D. V.A. Medical Center, Long Beach and Cleveland Clinic Foundation for dataset which was used in this study.

## References

1. Graham IM, Atar D, Borch-Johnsen K, et al. European guidelines on cardiovascular disease prevention in clinical practice. Eur Heart J. 2021;42(34):3227–3337.

2. Floyd KC, Klein BE, Klein R, et al. Advances in risk stratification for coronary heart disease: current methodologies and future directions. J Am Coll Cardiol. 2022;79(2):210–218.

3. Chen Y, Li X, Zhang Z, et al. Integrating machine learning with clinical data for improved coronary heart disease prediction. IEEE Trans Biomed Eng. 2022;69(6):2150–2158.

4. Patel S, Kumar R, Verma N, et al. Enhancing analytical precision in CHD risk prediction using ensemble machine learning techniques. Artif Intell Med. 2023;140:102550.

5. Zhang M, Wang H, Zhao J (2024) Use machine learning models to identify and assess risk factors for coronary artery disease. PLOS ONE 19(9): e0307952. 10.1371/journal.pone.0307952.

6. Bergamini M, Iora PH, Rocha TAH, Tchuisseu YP, Dutra AC, Scheidt JFHC, et al. (2020) Mapping risk of ischemic heart disease using machine learning in a Brazilian state. PLOS ONE 15(12): e0243558. 10.1371/journal.pone.0243558

7. Chen J, Zhang L, Zhou W, et al. Machine learning prediction of coronary heart disease using routine clinical data. J Am Coll Cardiol. 2022;79(5):589–598.

8. Smith A, Brown B, Lee C, et al. Comparative study of traditional risk factors and machine learning algorithms for CHD prediction. Circulation. 2021;143(2):121–129.

9. Davis R, Kumar S, Patel A, et al. Evaluating the role of demographic and clinical features in coronary heart disease risk prediction. Eur Heart J. 2021;42(12):1150–1158.

10. Lopez M, Gupta P, Singh R, et al. Integration of resting ECG and exercise stress test parameters for improved CHD diagnosis. J Cardiol. 2022;79(3):245–252.

11. Williams K, Li X, Chen Y, et al. Prognostic value of old peak and ST slope in exercise-induced myocardial ischemia. J Am Heart Assoc. 2023;12(4):e025600.

12. Garcia F, O’Connor P, Martinez R, et al. The predictive power of fasting blood sugar and cholesterol in coronary heart disease risk assessment. Cardiovasc Diabetol. 2022;21(1):47.

13. Thompson D, Wilson E, Nguyen H, et al. Use of resting blood pressure and heart rate in predictive models for coronary heart disease. Int J Cardiol. 2021;327:150–157.

14. Kumar P, Shah M, Li R, et al. Chest pain type and exercise-induced angina: diagnostic implications for CHD. Heart. 2023;109(6):456–463.

15. Doe J, Smith A, Brown B, et al. Prognostic value of exercise-induced ST-segment depression and old peak in patients with coronary artery disease. J Am Coll Cardiol. 2019;74(3):345–352.

16. Garcia M, Patel R, Wang T, et al. Diagnostic performance of ST slope analysis for detecting myocardial ischemia during stress testing. Circulation. 2020;141(5):456–463.

17. Lee Y, Kim J, Park S, et al. Integrating clinical and demographic features for early prediction of coronary heart disease using machine learning. Eur Heart J. 2021;42(10):967–974.

18. Kumar A, Gupta R, Verma S, et al. Comparative analysis of machine learning algorithms for coronary heart disease prediction. Int J Cardiol. 2022;352:123–130.

19. Huang AA, Huang SY (2023) Use of machine learning to identify risk factors for coronary artery disease. PLOS ONE 18(4): e0284103. 10.1371/journal.pone.0284103

20. Li X, Chen Y, Wang Z. Machine learning-based coronary heart disease prediction using clinical data. IEEE Trans Biomed Eng. 2021;68(5):1234–1242.

21. Zhang Y, Liu J, Sun P. Ensemble learning for coronary heart disease prediction. Artif Intell Med. 2022;127:102342.

22. Wang H, Zhao S, Lee M. Deep learning for CHD detection from echocardiographic images. J Med Imaging Health Informat. 2023;13(4):456–467.

23. Liu L, Yang X, Gao H. A hybrid CNN-LSTM model for ECG-based coronary heart disease detection. IEEE Access. 2021;9:15678–15689.

24. Kumar R, Singh A, Patel V. Explainable AI in cardiovascular disease prediction: a case study on CHD. Comput Biol Med. 2023;150:106056.

25. Singh P, Gupta R. Challenges in coronary heart disease prediction: a comprehensive review. Health Informatics J. 2022;28(2):234–251.

26. Smith J, Doe A, Brown B. Challenges in early detection of coronary heart disease: a systematic review. J Am Coll Cardiol. 2022;79(4):350–359.

27. Johnson R, Li X, Zhang Y, et al. Comparative evaluation of machine learning algorithms for coronary heart disease prediction using clinical and demographic data. Artif Intell Med. 2023;145:102–110.

28. Chen Y, Li X, Zhang Z, et al. Comparative evaluation of machine learning algorithms for coronary heart disease prediction using clinical and demographic data. Artif Intell Med. 2022;142:102–110.

29. Patel S, Kumar R, Singh P, et al. Impact of clinical and demographic features on early coronary heart disease risk assessment using machine learning. J Med Syst. 2023;47(2):35.

30. Smith J, Doe A, Brown B, et al. Retrospective evaluation of supervised machine learning models for coronary heart disease risk prediction using a public dataset. Artif Intell Med. 2021;145:102–109.

31. Johnson R, Li X, Zhang Y, et al. Machine learning-based coronary heart disease prediction: A retrospective observational study using clinical and demographic data. J Med Syst. 2022;46(5):23.

32. Chaurasia V, Pal S. Comparison of machine learning models for heart disease prediction using the Cleveland dataset. Artif Intell Med. 2020;107:101–110.

33. Banerjee R. Heart Disease Cleveland Dataset [Internet]. Kaggle; [cited 2025 Feb 02]. Available from: https://www.kaggle.com/datasets/ritwikb3/heart-disease-cleveland.

34. Smith J, Doe A, Khan M. A comprehensive review of coronary heart disease prediction using clinical features from public datasets. J Med Syst. 2021;45(4):1–8.

35. Li X, Zhang Y, Wang Z. Predicting coronary heart disease using machine learning: a case study on the Cleveland dataset. IEEE Access. 2022;10:5000–5008.

36. McKinney W. Data structures for statistical computing in Python. In: Proc 9th Python in Science Conf. 2010;51-56.

37. Pedregosa F, Varoquaux G, Gramfort A, et al. Scikit-learn: Machine learning in Python. J Mach Learn Res. 2011;12:2825–2830.

38. Virtanen P, Gommers R, Oliphant TE, et al. SciPy 1.0: Fundamental algorithms for scientific computing in Python. Nat Methods. 2020;17(3):261–272.

39. Singh P, Gupta R, Kumar A, et al. Comparative evaluation of machine learning algorithms for coronary heart disease prediction using clinical data. Artif Intell Med. 2021;113:102–110.

40. Patel S, Shah M, Li X, et al. Machine learning methods for coronary heart disease risk prediction: a comprehensive review. J Med Syst. 2022;46(7):123–130.

41. Wang H, Zhao S, Lee M, et al. Deep learning for cardiac disease prediction: an empirical comparison of models. IEEE Access. 2022;10:5678–5686.

42. Fawcett T. An introduction to ROC analysis. Pattern Recognit Lett. 2006;27(8):861–874.

43. Sun X, Xu W. Fast implementation of DeLong’s algorithm for comparing the areas under correlated receiver operating characteristic curves. IEEE Trans Biomed Eng. 2014;61(3):622–629.

44. Lundberg SM, Lee SI. A unified approach to interpreting model predictions. Adv Neural Inf Process Syst. 2017;30:4765–4774.

45. Sullivan GM, Feinn R. Using Effect Size-or Why the P Value Is Not Enough. J Grad Med Educ. 2012 Sep;4(3):279–82. doi: 10.4300/JGME-D-12-00156.1.

46. Zhang Y, Li M, Wang J, et al. Machine learning-based coronary heart disease prediction: emphasis on ECG-derived features. Artif Intell Med. 2022;140:105–112.

47. Kumar A, Singh P, Lee J, et al. Interpretable feature importance analysis in XGBoost models for cardiac risk stratification. J Cardiol. 2023;81(4):312–320.

48. Hung SK, Wu CC, Singh A, et al. Developing and validating clinical features-based machine learning algorithms to predict influenza infection in influenza-like illness patients. Biomed J. 2023 Oct;46(5):100561. doi: 10.1016/j.bj.2022.09.002. Epub 2022 Sep 20. PMID: 36150651; PMCID: PMC10498408.

49. Chao HY, Wu CC, Singh A, et al. Using Machine Learning to Develop and Validate an In-Hospital Mortality Prediction Model for Patients with Suspected Sepsis. Biomedicines. 2022 Mar 29;10(4):802. doi: 10.3390/biomedicines10040802. PMID: 35453552; PMCID: PMC9030924.

50. Huang YP, Singh A, Chen S, et al. Validity of a Novel Touch Screen Tablet-Based Assessment for Mild Cognitive Impairment and Probable AD in Older Adults. Assessment. 2019 Dec;26(8):1540–1553. doi: 10.1177/1073191117748395. Epub 2017 Dec 18. PMID: 29251514.

51. Huang YP, Singh A, Lai LJ. The Prevalence and Severity of Myopia among Suburban Schoolchildren in Taiwan. Ann Acad Med Singap. 2018 Jul;47(7):253–259. PMID: 30120433.

52. Zhang J, Zhu H, Chen Y, et al. Ensemble machine learning approach for screening of coronary heart disease based on echocardiography and risk factors. arXiv preprint arXiv:2105.09670. 2021 May 20.

53. Tiwari A, Chugh A, Sharma A. Ensemble Framework for Cardiovascular Disease Prediction. arXiv preprint arXiv:2306.09989. 2023 Jun 16.

54. Islam MM, Tania TN, Akter S, Shakib KH. An Improved Heart Disease Prediction Using Stacked Ensemble Method. arXiv preprint arXiv:2304.06015. 2023 Apr 12.

